# Time of Day Dependent Effects of Contractile Activity on the Phase of the Skeletal Muscle Clock

**DOI:** 10.1101/2020.03.05.978759

**Authors:** Denise Kemler, Christopher A. Wolff, Karyn A. Esser

## Abstract

Exercise has been proposed to be a zeitgeber for the muscle circadian clock mechanism. However, this is not well defined and it is unknown if exercise timing induces directional shifts of the muscle clock. Our purpose herein was to assess the effect of one bout of treadmill exercise on skeletal muscle clock phase changes. We subjected PERIOD2::LUCIFERASE mice (n=30F) to one 60-minute treadmill exercise bout at three times of day. Exercise at ZT5, 5h after lights on, induced a phase advance (1.4±0.53h; p=0.038), whereas exercise at ZT11, 1h before lights off, induced a phase delay (−0.95±0.43h; p=0.0315). Exercise at ZT17, middle of the dark phase, did not alter muscle clock phase. Exercise induces diverse systemic changes so we developed an *in-vitro* model system to examine effects of contractile activity on muscle clock phase. Contractions applied at peak or trough *Bmal1* expression induced significant phase delays (applied at peak: 1.3±0.02h; p=0.0425; applied at trough: 1.8±0.02h, p=0.0074). Contractions applied during the transition from peak to trough *Bmal1* expression induced a phase advance (1.8±0.03h; p=0.0265). Lastly, contractions at different times of day resulted in differential changes of core-clock gene expression demonstrating an exercise and clock interaction, providing insight into potential mechanisms exercise-induced phase shifts. These data demonstrate that muscle contractions, as part of exercise, are sufficient to shift muscle circadian clock phase, likely through changes in core-clock gene expression. Additionally, our findings that exercise induces directional muscle clock phase changes confirms exercise is a bone fide environmental time cue for skeletal muscle.

## INTRODUCTION

The circadian clock is an evolutionarily conserved regulatory mechanism that allows organisms to adapt, respond, and entrain to their environment. By integrating/responding to environmental signals (i.e., time cues), the circadian clock coordinates daily oscillations of behaviour, metabolism, and gene expression at the organismal and tissue-specific levels driving circadian rhythms (Golombek and Rosenstein, 2010). Contained in virtually all cells throughout the body, the mammalian circadian clock is an endogenous self-sustaining transcription-translation feedback loop that directs a daily program of gene expression. Consistent across tissues, the core clock transcription factors *Bmal1* and *Clock* drive the expression of *Period1/2* and *Cryptochrome 1/2* which in turn repress their own expression by inhibiting BMAL1 and CLOCK transcriptional activity (Ko and Takahashi, 2006; McCarthy et al., 2007; Mohawk et al., 2012; Partch et al., 2014). The circadian cycle takes ≈24h to be completed and thus defines the period length (e.g. interval between two peaks) of the circadian clock output.

While the period length is fixed, the circadian phase is variable and represents a temporal reference point relative to a fixed event. The most prominent example is the central clock in the brain (SCN) where the circadian phase of a rest/activity pattern is determined by the naturally occurring light/dark cycle (fixed event) driving most circadian behaviour (Merrow et al., 2005; Wright et al., 2013). Environmental time cues that are capable of entrainment (i.e., setting circadian phase) are called Zeitgebers and while light is the most prominent time cue, it primarily entrains the SCN central clock. However, additional non-photic zeitgebers exist such as feeding, activity, and stress, all of which serve as time cues to peripheral tissue clocks (Fuller et al., 2008; Mistlberger and Antle, 2011; Sujino et al., 2012; Tahara and Shibata, 2018). Therefore, light acts as a zeitgeber for the central clock, setting a phase for all clocks throughout the body. However, non-photic time cues can act on peripheral tissues, influencing the phase of peripheral tissue clocks with the potential for putting them out of alignment with the phase set by the SCN (Koronowski et al., 2019; Maywood and Mrosovsky, 2001; van der Vinne et al., 2018; Wolff and Esser, 2012).

Time cues must meet certain criteria to qualify as a true zeitgeber. Specifically, an environmental time cue must interact with the core molecular clock mechanism to shift the phase of the endogenous rhythm. This shift in phase can be a negative (delay) or positive (advance) and its direction is dependent on the time the environmental cue was applied. Finally, the phase shift must be persistent in contrast to a local reset in which case the phase returns to its old status before the time cue was applied. A powerful tool to investigate the effects of zeitgeber on the circadian phase is the phase response curve (PRC). Phase response curves are a graphic representation of the dynamics (phase shift direction and magnitude) of a circadian oscillator when stimulated by a zeitgeber at different times of day (reviewed in Johnson, 1999). When a PRC shows a phase shift during the subjective night (usually advance in the first half and a delay in the second) they are called “photic-PRC” (Daan and Pittendrigh, 1976; Rusak and Boulos, 1981). Those curves with phases that advance during the dark phase (are so called “non-photic” PRC and happen in response to stimuli different from light (Eastman et al., 1995a; Mistlberger and Antle, 2011; Mrosovsky, 1996; Reebs and Mrosovsky, 1989; Youngstedt et al., 2019). Thus, PRCs provide insight into whether a cue is a zeitgeber and how it modulates the phase of a circadian oscillator e.g. tissue specific clocks.

Exercise is a robust stimulus that induces significant changes in core temperature, and hormonal responses as well as tissue specific transcriptional and metabolic outcomes. Skeletal muscle is a large organ system and is known to be an important tissue for overall organismal health (Demontis et al., 2013). Exercise exerts numerous positive effects on skeletal muscle health (Cartee et al., 2016; Gabriel and Zierath, 2019) and proper muscle circadian function is important to maintain skeletal muscle health (Chatterjee and Ma, 2016; Dyar et al., 2018; Harfmann et al., 2016; Kondratov et al., 2006; Schiaffino et al., 2016; Schroder et al., 2015). To date, however, it is unclear how exercise interacts with the muscle clock and if exercise is a bona fide zeitgeber for the circadian clock in skeletal muscle. In the current investigation, we use a combination of an established mouse model of exercise and a newly designed in-vitro cell system to demonstrate that muscle contractions as a component of exercise are indeed a zeitgeber for the circadian clock in skeletal muscle. Moreover, we also investigate how contractions modify the phase of the muscle circadian clock as well as expression of core clock genes independent of external circadian inputs. These observations will lay the foundation for further research to interrogate the connection between exercise and the muscle clock to better understand the potential for use of time of day exercise interventions for therapeutic effects.

## Methods

### Ethical Approval

All animal procedures in this study were conducted in accordance with the guidelines of University of Florida for the care and use of laboratory animals (IACUC #201809136). The use of animals for exercise protocols were in accordance with guidelines established by the U.S. Public Health Service Policy on Humane Care and Use of Laboratory Animals.

### Treadmill Exercise in PERIOD2::LUCIFERASE Mice

We used 30 female PERIOD2::LUCIFERASE (PER2::LUC)(Yoo et al., 2004) mice (3mo: n=20 20±1g; 14mo: n=10 25±2g) to test the effects of an acute bout of treadmill exercise on the circadian clock in skeletal muscle. Mice were group housed (2-5/cage) and maintained on a 12:12h light:dark schedule with ad libitum access to standard rodent chow (Envigo Teklad 2918, Indianapolis, IN, USA) and water. Briefly, following two days of treadmill acclimation at ZT5, ZT11, or ZT17 (for reference ZT0 = lights on: ZT12 = lights off: ZT17 was completed under red light) (Figure 1A) five PER2::LUC animals were subjected to 1h of treadmill running at 15m/min at 0° (EX) on a Panlab treadmill (Harvard Apparatus, Holliston, MA) while five other PER2::LUC animals were moved from the housing suite into the treadmill room and were maintained sedentary for the duration of the exercise (Figure 1A). Food and water were removed from both groups at the onset of exercise and not returned. Following the cessation of exercise, mice were returned to their home cages for 1h. Mice were anesthetized using isoflurane and euthanized by cervical dislocation. EDL muscles were dissected from the mouse, taking care to isolate from tendon to tendon and used for real time bioluminescence recording.

**Figure 1:**
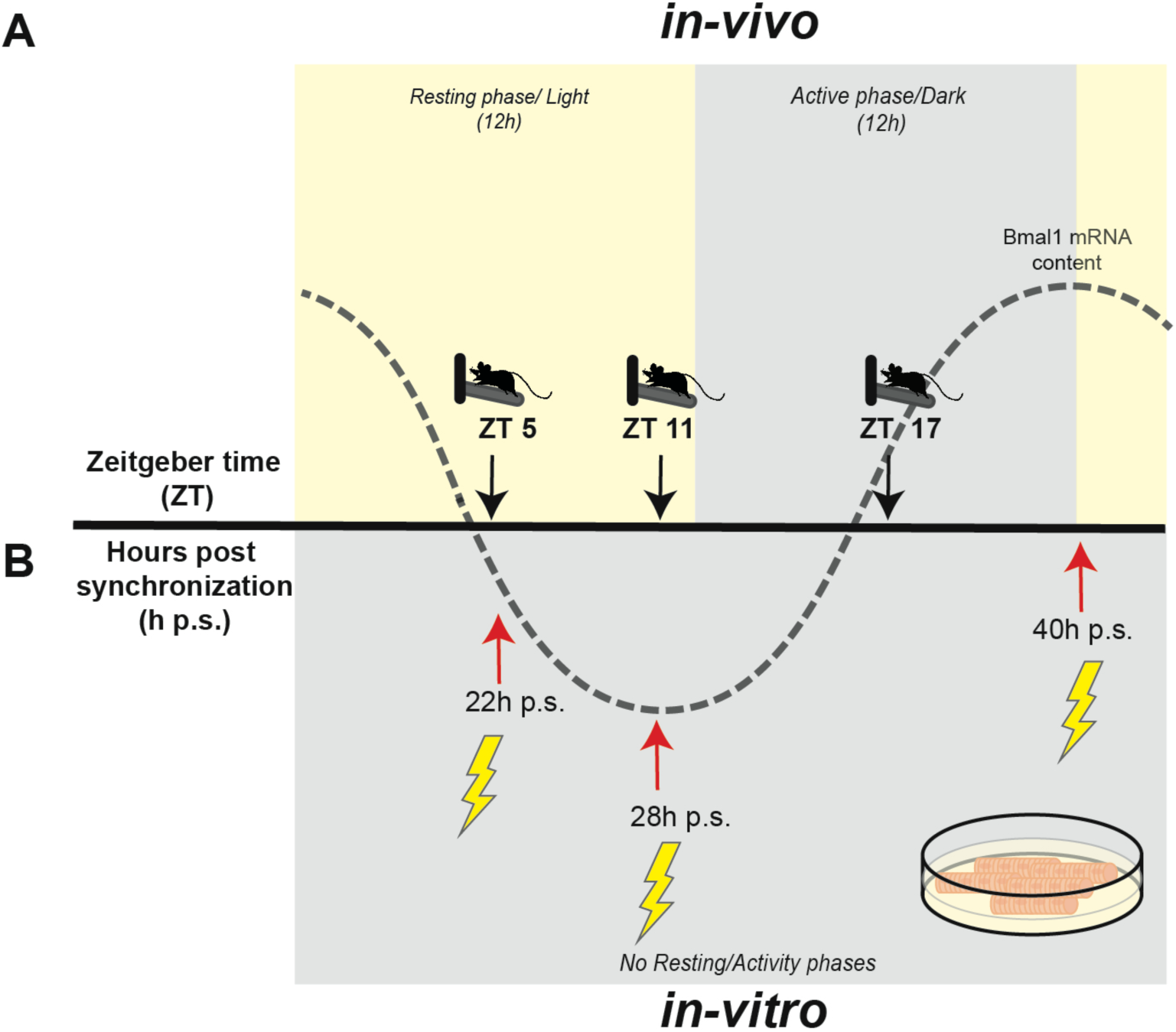
Timing of exercise/contractions in mice and in myotubes in cell culture. (A) Mice were exercised for 1h in the middle or end of the resting phase (ZT5 or ZT11) and in the middle of the active phase (ZT17). Timing of exercise is defined as Zeitgeber Time (ZT), which represents time based on the period of a Zeitgeber (time cue) in this case 12h light:12h dark. (B) C2C12 myotubes were synchronized with dexamethasone and electrical pulse stimulated for 1h to induce contractions (lightning bolt). The timing was aligned along the Bmal1 mRNA content in the cells and is presented in hours post synchronization (h p.s.).

### Cell Culture, Myogenic Differentiation and Circadian Synchronization

C2C12 cells were purchased from ATCC (ATCC Cat# CRL-3419, RRID:CVCL_UR38, LOT: 70004012). Cells were cultured CO_2_, 95% humidity at 37°C. Cells were maintained in 150mm cell culture dishes in growth media (GM) consisting of Dulbecco’s Modified Eagle’s Medium (DMEM) supplemented with 10% fetal bovine serum (FBS) and 1% penicillin-streptomycin (P/S) and 1mM Sodium Pyruvate. C2C12 cells were grown to 100% confluency on 35mm cell culture dishes that were previously coated with a gelatine hydrogel as published before with some variations to the protocol (Bettadapur et al., 2016; Denes et al., 2019). The non-patterned hydrogel was made by dissolving 12%w/v porcine gelatine (Sigma, St. Louis, MO) in ddH_2_O by heating to 65°C and dissolving 10% w/v solution of microbial transglutaminase (MTG) by heating to 37°C. Gelatine and MTG were mixed 10:1 and incubated for 5min at 65°C. 120 µl of gelatine was added to the 35mm dishes, distributed equally, and crosslinked at room temperature for 16h-18h. Gelatine coated plates were rehydrated with phosphate buffered saline (PBS) for 4h and UV sterilized for 30min prior to use. For cell differentiation, medium was changed to differentiation medium (DM) consisting of DMEM supplemented with 2% horse serum (HS) and 1% P/S + 1mM of Sodium Pyruvate. Cells were differentiated for 3 days before experiments were performed. For synchronization of circadian clocks myotubes were treated with 1µM dexamethasone (Sigma-Aldrich D2915) in DM for 90 minutes (Balsalobre et al., 2000). After treatment cells were washed with PBS and received fresh DM.

### Generation Of Stable Transfected C2C12 Cells

Stable C2C12 cells were generated using non-viral DNA transposon (Balciunas et al., 2006a). The Bmal1P-Luc reporter gene contains the wildtype Bmal1 promoter sequence starting 394 bp upstream of the transcription start (TSS) and ending 154 bp downstream of the TSS and was a gift from Dr. John Hogenesch (Sato et al., 2004). Bmal1P-Luc was subcloned into a modified version of pminiTol2 plasmid (removal of CMV promoter and addition of pSV40neo cassette), a gift from Stephen Ekker (Addgene plasmid # 31829; http://n2t.net/addgene:31829; RRID:Addgene_31829). The transposase pCMV-Tol2 was a gift from Stephen Ekker (Addgene plasmid # 31823; http://n2t.net/addgene:31823; RRID:Addgene_31823; Balciunas et al., 2006b). Transfected cells were selected for integration of the transgene over the course of one week with G418 (2mg/ml). Clones that tested positive for transgene integration and that showed sustained bioluminescence oscillations with a period length of 23-24h were further cultivated. For the final experiments three different clones were used to generate the data for each time point to exclude that observed results were influenced by the transgene integration into the genome.

### Electrical Pulse Stimulation Of C2C12 Myotubes

Day three, C2C12 myotubes received fresh DM 1-3 hours prior to electrical pulse stimulation (EPS). EPS was carried out in 35mm dishes using IONOPTIX C-Pace EP system, stimulating 6 dishes in the same bank. Conditions were 25ms pulse with 10V at 6Hz every 5 seconds for one-hour, visual assessment of all plates revealed twitching across the field of view. Each EPS experiment included 3-5 technical replicates per treatment group (no-stim vs. EPS), and each new EPS experiment was conducted using a distinct biological replicate of cells, including different clones of Bmal-luc stable cell lines. Myotubes were stimulated at 22h, 28h, or 40h post synchronization (Figure 1B), placed into recording medium, vacuum sealed and put into the Lumicycle for bioluminescence recording.

### RNA Isolation and Quantitative Real-Time PCR

Three dishes (35mm) were collected in 500µl of Trizol each (Invitrogen 15596018) and then pooled into one tube and were treated as a single biological replicate. RNA was isolated using the RNeasy Mini Kit (Qiagen 74104) according to the manufacturer’s protocol. DNAse digest was performed on column using the RNase-Free DNase Set (Qiagen 79254) according to manufacturer’s protocol. cDNA was generated using 500ng of total RNA using SuperScript III First Strand Synthesis System (Thermo Fisher Scientific 18080051) according to manufacturer’s protocol. All cDNA samples were diluted 1:25 in RNAse free water and 4µl were used to perform quantitative real time PCR (qRT-PCR). The qRT-PCR was carried out using the Applied Biosystems Fast SYBR Green Master Mix (Thermo Fisher Scientific 43-856-14) with 10µM of each primer. Primer sequences are Bmal1for TCAAGACGACATAGGACACCT, Bmal1rev GGACATTGGCTAAAACAACAGTG; Per1for CGGATTGTCTATATTTCGGAGCA, Per1rev TGGGCAGTCGAGATGGTGTA; Per2for AAAGCTGACGCACACAAAGAA, Per2Rev ACTCCTCATTAGCCTTCACCT; Cry1for CACTGGTTCCGAAAGGGACTC, Cry1rev CTGAAGCAAAAATCGCCACCT; Cry2for CACTGGTTCCGCAAAGGACTA, Cry2rev CCACGGGTCGAGGATGTAGA; RevErbαfor TCCCAGGCTTCCGTGACCTTT, RevErbα TTGTGCGGCTCAGGAACATCA. qRT-PCR was performed in a QuantiStudio 3 thermal cycler (Applied Biosystems, Foster City, CA). mRNA levels of target genes were normalized using *Rpl26* mRNA levels and relative quantification was calculated by using the ΔΔct method. We used Rpl26 for normalization as it does not exhibit an oscillatory expression pattern (CircaDB) and is not responsive to EPS. To determine if the expression of a given mRNA exhibited a circadian oscillation, we utilized a Single Cosinor analysis (Refinetti et al., 2007). In order to measure the effect of EPS on circadian clock gene expression, mRNA levels of target genes were normalized using *Rpl26* mRNA (Rpl26for CGAGTCCAGCGAGAGAAGG, Rpl26rev GCAGTCTTTAATGAAAGCCGTG) levels and relative quantification was calculated by using the ΔΔct method. Data for each independent experimental replicate is presented as the fold change from control (ΔΔCT EPS/ΔΔCT Con). We present the mRNA expression data as the EPS-induced percent change relative to Control.

### Real Time Bioluminescence Recordings

For real time bioluminescence recording, myotubes were washed with PBS after the treatment and switched to recording medium: DMEM without phenol red (Caisson DML12-500ml) supplemented with 2% HS, 1mM Sodium Pyruvate, 1%P/S and 0.1mM Luciferin. Tissue explants were cultured in a similar recording medium using 5% FBS in place of 2% HS. Dishes were vacuum sealed with a microscopy glass coverslip and placed into the Lumicycle 32 (Actimetrics, Wilmette, IL). Real time bioluminescence recording was performed with a sampling frequency of every 10 minutes for at least four consecutive days as previously described in detail by our laboratory (Wolff and Esser, 2012). We removed the first 24h of baseline-subtracted raw data, due to the expected measurement fluctuations within the first 24h of recording. Trimmed data were analysed using JTK_Cycle (RRID:SCR_017962) to determine Phase (lag), Period length and Amplitude. Benjamini-Hochberg multiple comparisons adjusted p-value was used to assess the quality of the curve fit. For the cell culture myotube experiments we included at least three biological replicates for each time point and three to five technical replicates for each biological replicate group. No *in vitro* data were excluded. For the PER2::LUC tissue explant experiments, each extensor digitorum longus (EDL) was considered a technical replicate for a given animal. Animals were excluded whose explanted muscles did not show any circadian oscillation (n=1 at ZT5) or technical replicates (n=1 at ZT11 and n=2 at ZT17) were remove, when noisy readings interfered with calculation of circadian phase. One muscle ripped during preparation and could not be used (ZT5, EDL5-2).

### Statistical Analyses

Statistical analyses were completed using GraphPad Prism 8.3 (GraphPad Prism, RRID:SCR_002798). For Cosinor Analysis a curve was fitted using a predefined period of 24h by the least square’s method. Rhythm characteristics including 95% confidence interval include the MESOR (middle value of the fitted cosine representing a rhythm adjusted mean), amplitude (half the difference between the minimum and maximum of the fitted cosine function) and the acrophase (peak of a rhythm). Significance of rhythm was determined by rejection of the zero-amplitude hypothesis with a threshold of 95%. Pooled correlation coefficient values from each biological replicate (n=3 independent experiments) were used to calculate mean, standard deviation, and sample size for a one-way ANOVA to examine changes in Cosinor goodness of fit for the duration of the time course experiments.

For analysis of summarized luminescence data, as well as qPCR data we were *a priori* interested in the effect of EPS on circadian phase and core clock gene expression independent of time of day, and thus compared the EPS-induced change in mRNA expression or circadian phase to untreated control samples using an unpaired Student’s t-tests once we confirmed data were normally distributed using Shapiro-Wilks Normality Tests. Differences between groups were considered significant at the level of p<0.05. All data are presented as Mean±SD.

## RESULTS

### Time-Specific Effects of Exercise on Circadian Phase in EDL

Previous data from our group found that four weeks of daily treadmill running in the rest/light phase resulted in significant shifts of the skeletal muscle clock, with no shift in the central clock in the SCN (Wolff and Esser, 2012). We sought to determine if a single 60-minute bout of moderate intensity treadmill running was sufficient to alter the phase of muscle clocks. We also assessed the effects of exercise timing relative to rest/activity on muscle circadian rhythms to find out if exercise functions as a zeitgeber capable of inducing both phase advances and phase delays of the muscle clock.

PER2::LUC mice completed a one-hour moderate intensity exercise bout at either ZT5 (middle of the light/resting phase) ZT11 (end of light/resting phase) or ZT17 (middle of the dark/active phase). EDL muscles showed PER2::LUC oscillation in control (black) and exercised (blue) cohorts and this oscillation was maintained over several days for all three time points (Figure 2A and B). We analysed for changes in phase by calculating the phase of control vs. exercised mice using JTK cycle, and calculating the fold change of stimulated vs unstimulated. We found that exercise at either ZT5 or ZT11 resulted in significant changes in the phase of bioluminescence that this change in phase was maintained over the full course of the experiment (Figure 2A). Exercise at ZT5, the middle of the rest phase, induced a significant (p= 0.0387) phase advance of 1.4±0.53h (Figure 2B left) whereas exercise at ZT11, at the end of the rest phase, induced a significant (p=0.0315) phase delay of −0.95±0.40h (Figure 2B middle). In contrast to exercise at ZT5 and ZT11, exercise at ZT17, in the middle of the dark/active phase did not significantly shift (p=0.4808) the phase of the muscle circadian clock (−0.67±1.1h: Figure 2B right). Taken together, our findings demonstrate that an acute bout of treadmill running is sufficient to functions as a time cue for the muscle clock causing differential shifts depending on time of exercise. We also note that one bout of treadmill exercise did not result in changes in the amplitude of period length of the EDL muscle clock (data not shown).

**Figure 2.**
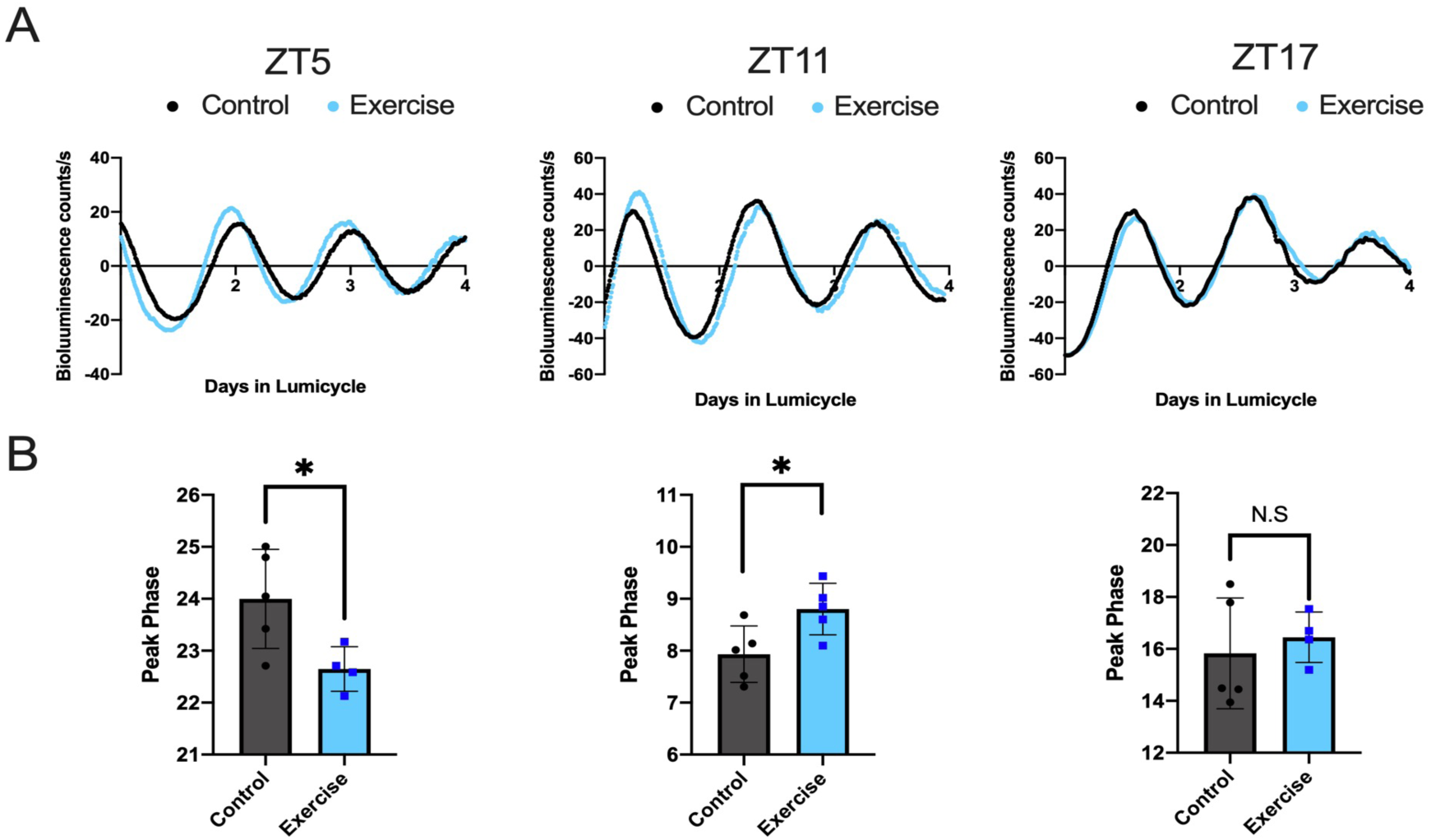
One Hour of Treadmill Running at Different Times During the Circadian Cycle Shifts Phase of the Muscle Clock Differentially. (A) Bioluminescent tracing (baseline subtracted) of PER2::LUC activity in explanted EDL muscles after 1h of acute treadmill run exercise over 4 days (mean of n=4 biological replicates) at three different time points ZT5, ZT11 or ZT17. Control animals in black, exercise group in blue. (B) Effect of acute treadmill running exercise at ZT5, ZT11 or ZT17 on the peak phase of EDL muscle PER2::LUC bioluminescence (n= 5 biological replicates). Circadian phase was calculated using time of peak luminescence after 24h in the Lumicycle utilizing JTK cycle. **ZT5** p= 0.0387, **ZT11** p= 0.0315, **ZT17** p=0.4808 (nested T-test). Decrease in peak phase = phase advance, increase in peak phase = phase delay.

### C2C12 myotubes exhibit an endogenous circadian rhythm

We demonstrate that a single bout of running exercise is sufficient to alter the phase of the muscle clock in a time-specific manner. However, running is a systemic physiological stimulus that is associated with hormonal changes, core temperature changes as well as local muscle metabolism. Thus, our next set of experiments was designed to test the role of muscle contractions as a time cue for the clock. To investigate the effect of contractions on the circadian clock in skeletal muscle in an isolated system, we established an in-vitro model, with a single cell type. C2C12 myotubes are commonly used to study various aspects of skeletal muscle physiology. In order to serve as a model for our purposes, we first ran time course studies to validate that the endogenous clock factors oscillate over time in C2C12 myotubes. We synchronized the phase of C2C12 myotubes with brief exposure to dexamethasone then analysed the mRNA expression of seven core clock genes (*Bmal1, Clock, Per1, Per2, Cry1, Cry2, RevErbα*) as well as *Dbp*, an established clock-output gene, every 4h over 24h (Figure 3). Using Cosinor analysis we determined that *Bmal1 (p=0*.*036), Per1 (p=0*.*0511), Per2 (p=0*.*0356), Cry1 (p=0*.*0057)*, and *Dbp (p=0*.*002)* exhibited significant circadian expression, but *Cry2 (p=0*.*9693)* and *Clock (p=0*.*3710)* did not. Rev-Erbα did not reach statistical significance (p=0.0723), as one replicate from an individual experiment at the 36h post synchronization time point was not in line with the others (black circled dot). We also determined that the peak of *Bmal1* mRNA expression occurred at ∼40h post synchronization, while the peak of *Per1/2* expression occurred at ∼28h, antiphase to *Bmal1*, as has been shown in mouse and human muscle circadian transcriptomes in the (McCarthy et al., 2007; Perrin et al., 2018; Pizarro et al., 2012).

**Figure 3.**
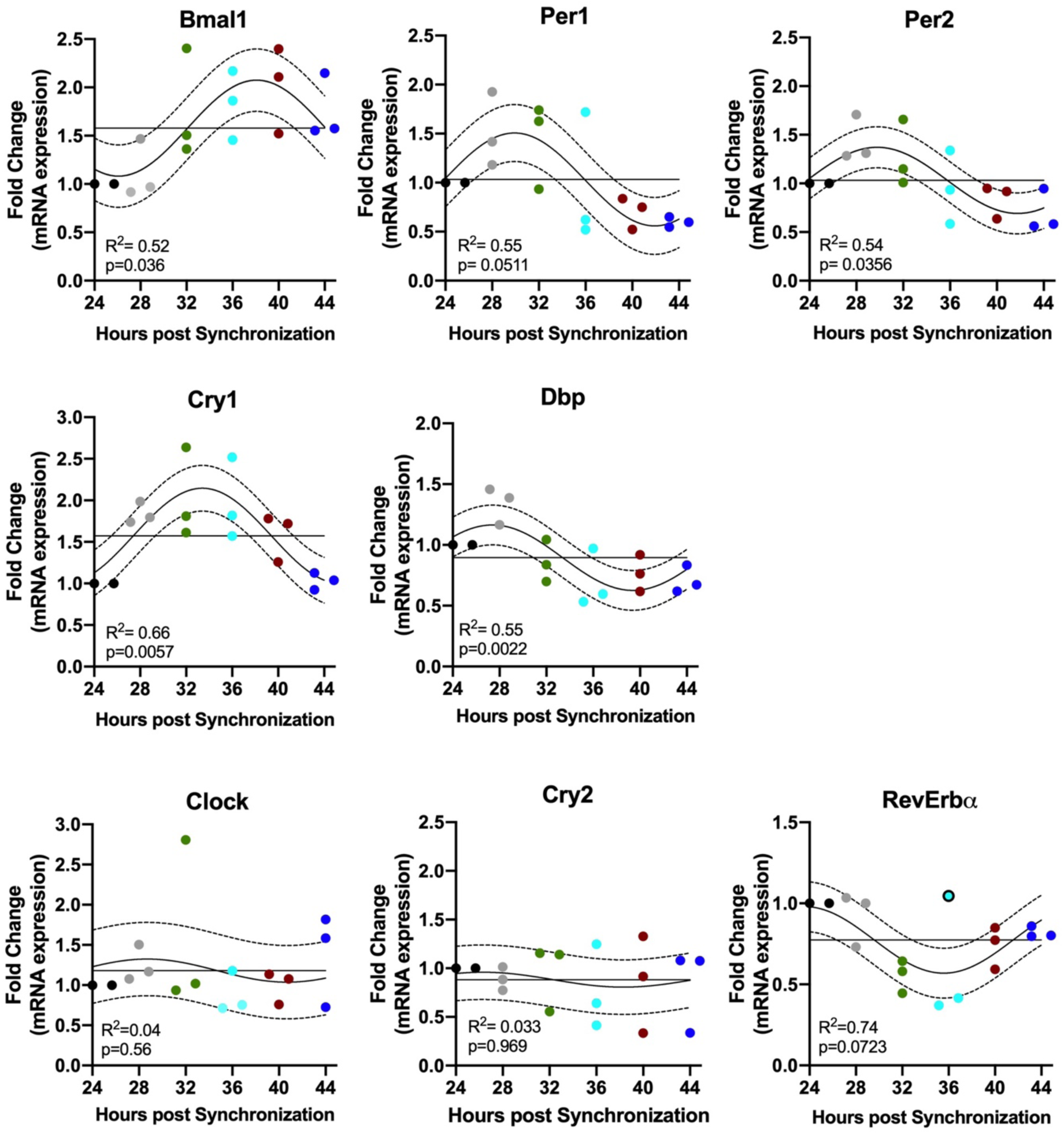
C2C12 Cells Exhibit an Endogenous Rhythm of Core Clock mRNA Expression. Twenty-four-hour gene expression profiles of known molecular clock-related genes in C2C12 myotubes. Gene expression levels are represented as fold change and normalized to the initial time point (24h). Individual values are displayed for n=3 biological replicate experiments at each different time point (depicted by different colours 24h: black; 28h: grey; 32h green, 36h light blue; 40h red; 44h dark blue). Each biological replicate for each time point is pooled from 3 technical replicates (i.e., 35mm dishes). To improve visibility of individual experimental replicates, time points where all data points were not visible were adjusted horizontally (e.g., Bmal1 28h). The best fitting 24h single cosine curve is shown. Dotted line represents the 95% confidence interval and the straight horizontal line the MESOR. Goodness of fit and the result of analysis of time of day variance are represented as R^2^ and p-values, respectively.

### One bout of electrical field stimulation changes the circadian phase in C2C12 myotubes

The next question we pursued was whether a single 60-minute bout of contractions is sufficient to change the phase of the skeletal muscle clock (e.g. Bmal1-luc) in the absence of systemic factors. We generated 20 stable C2C12 clones with a circadian clock reporter gene, Bmal1:Luc and for these experiments we used three separate clones to improve the rigor of this approach. We stimulated synchronized C2C12 myotubes (day 4 post differentiation) for one hour at either 22h following synchronization (22h p.s.), 28h following synchronization (28h p.s.), and 40h following synchronization (40h p.s.). These time points were chosen based on previously defined peak/trough expression level of Bmal1 mRNA in the cells (see Figure 1B).

Following stimulation, the dishes were placed in the Lumicycle and phase of the Bmal1-luc was assessed over multiple days measuring bioluminescence signal under static conditions (no medium change) from the Bmal1-Luc reporter. The myotubes exhibit a clear oscillatory pattern of bioluminescence over multiple days, confirming that our cell lines expressed the integrated Bmal1:Luc transgene in proper rhythmic fashion (Figure 4A). We analysed for changes in phase by calculating the phase of control myotubes vs stimulated myotubes using JTK cycle, and calculating the fold change of stimulated vs unstimulated. Stimulation at 22h p.s. lead to a significant phase advance of 1.8±0.03h (p< 0.0001) (Figure 4B left). In contrast, stimulation at 28h p.s. lead to a significant phase delay of 1.8±0.02h (p=0.0074) (Figure 4B middle), as did stimulation at 40h p.s. (1.3±0.02h; p=0.0425) (Figure 4B right). These results show that contractions by themselves are sufficient to function as a time cue for the skeletal muscle clock, without additional input from other cell types or tissues.

**Figure 4:**
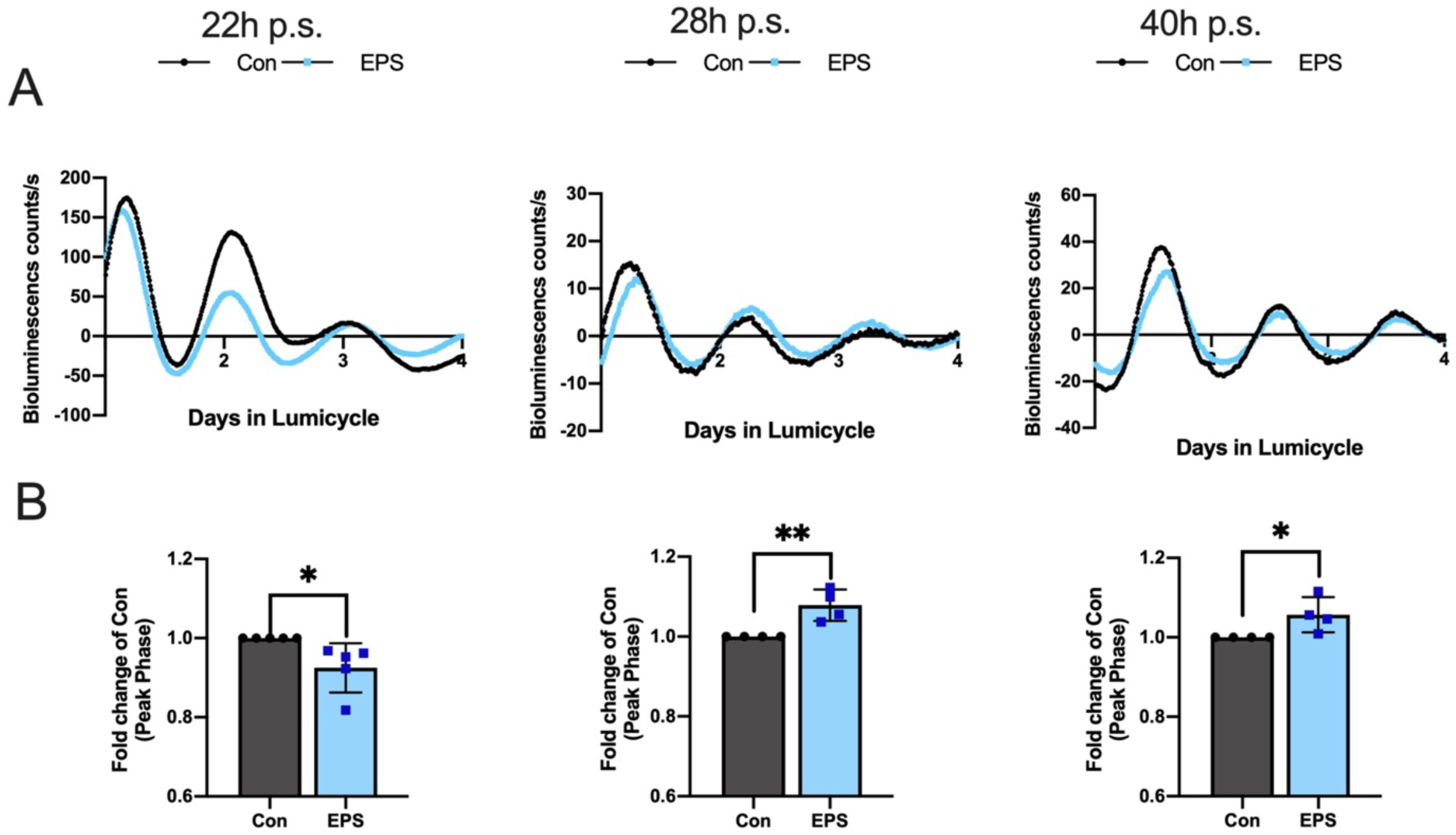
One bout of Contractions Phase Shifts the Clock in C2C12 Myotubes. (A) Bioluminescent tracing (baseline subtracted) of Bmal1:Luc activity in C2C12 myotubes after 1h of contraction inducing stimulation recorded over 4 days (representative tracing from one experiment, n=5 technical replicates) at three different time points 22h post synchronization (p.s.), 28h post synchronization (p.s.) or 40h post synchronization (p.s.). Control dishes in black, exercise group in blue. (B) Effect of contraction inducing stimulation three different time points 22h post synchronization (p.s.), 28h post synchronization (p.s.) or 40h post synchronization (p.s.). on the peak phase C2C12 myotubes, presented as Fold change of control Bmal1:Luc bioluminescence (n= 4 biological replicates). Circadian phase was calculated using the time of peak luminescence after 24h in the Lumicycle utilizing JTK cycle. **22h p**.**s**. p< 0.0001, **28h p**.**s**. p= 0.0074, **40h p**.**s**. p=0.0425 (T-test). Decrease in peak phase = phase advance, increase in peak phase = phase delay.

### Time-Specific Effects of Contraction on Molecular Clock Gene Expression

We have shown that contractions induce phase shifts of different directionality depending on time of day. This suggests that they might change the molecular clock gene program differently depending on time of exercise. For that reason, we investigated whether acute contractions lead to changes in gene expression in the positive arm components of the circadian clock (*Bmal1/Clock*) or the negative arm of the circadian clock (*Per1/2, Cry1/2*). For this set of experiments, we stimulated C2C12 myotubes under the same conditions and collected the cells for mRNA analysis. We analyzed expression of the core clock factors immediately post contractions. We found that contractions applied at 22h p.s. significantly (p=0.0150) reduced *Per2* mRNA expression compared to unstimulated controls by approximately 40%, but had no effect on *Per1* or *Bmal1* (Figure 5 left). However, when myotubes were stimulated at 28h p.s., we found that mRNA levels for *Bmal1* (p=0.0011), *Per1* (p=0.0197) and *Per2* (p<0.0001) were significantly decreased around 40% compared to unstimulated controls (Figure 5 middle) and no significant changes in any other core clock gene. Finally, unlike both other time points, myotubes stimulates at 40h p.s. did not significantly influence the expression of any core clock genes compared to unstimulated controls (Figure 5 right). These results show that a single bout of acute contractions is sufficient to induce changes in the steady state mRNA levels of key components of the circadian clock in a time-dependent manner.

**Figure 5.**
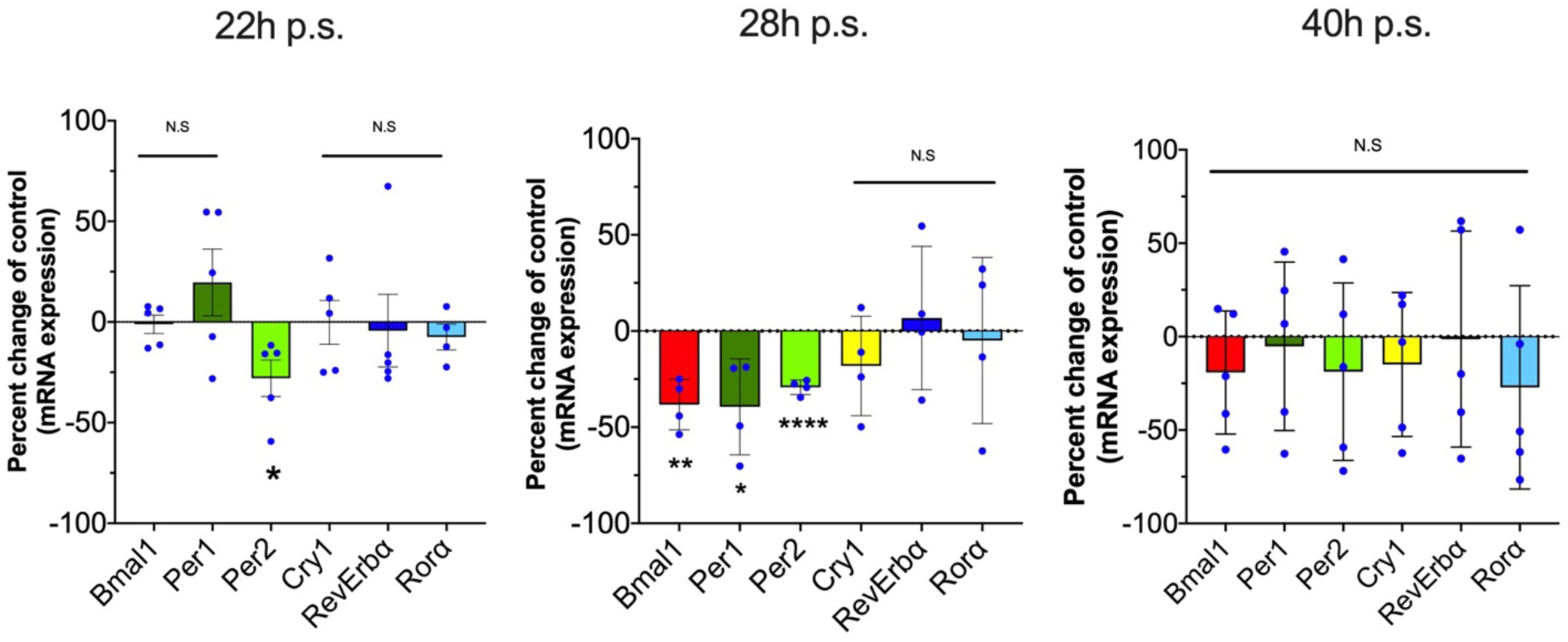
Contractions at Different Times During the Circadian Phase Modify Expression of Core Clock Genes Differentially. C2C12 were subjected to electrical field stimulation either at three different time points 22h post synchronization (p.s.), 28h post synchronization (p.s.) or 40h post synchronization (p.s.). Gene expression patterns of the core clock genes *Bmal1 (red), Per1(dark green), Per2 (light green), Cry1 (yellow), RevErbα(dark blue) and Rorα (light blue)* measured immediately post exercise. Data are presented as percent change of mRNA expression difference of EPS relative to control for each gene after normalization to *Rpl26* at each time point n= 5 biological replicates (22h/28h p.s.) n=4 biological replicates (28h p.s.) ± S.D. **22h p**.**s**. Per2 p= 0.0150; **28h p**.**s**. Bmal1 p= 0.0011, Per1= 0.0197, Per2 < 0.0001**; 40h p**.**s**. p > 0.2 for all genes (t-test).

## DISCUSSION

Exercise is well known stimulus that can alter skeletal muscle and organismal physiology. While emerging data has shown potent time-of-day specific outcomes of exercise on skeletal muscle gene expression and metabolism (Ezagouri and Asher, 2018; Gabriel and Zierath, 2019; Sato et al., 2019) the studies of the potential role of exercise as an environmental time cue for circadian clocks have been limited. Specifically studies have performed time of day exercise interventions and measured both behavioral as well as clock outcomes but all these studies performed repeated exercise training (Edgar and Dement, 1991; Marchant and Mistlberger, 1996; Wolff and Esser, 2012; Yamanaka et al., 2008). We report, for the first time, that a single one-hour bout of moderate intensity treadmill exercise is sufficient to differentially shift the phase of the circadian clock in skeletal muscle in mice. We next used an *in-vitro* model system, to remove the influence of systemic factors, to test the contribution of contractions to the phase shift of the muscle clock. We found that electrically induced contractions of muscle cells shift the phase of the circadian clock. The magnitude and direction of the time-specific effects of contractions on phase shifts were similar in vivo and in vitro models. Lastly, we found that the timing of contractions differentially altered the mRNA expression of core clock genes, *Per1, Per2*, and *Bmal1* providing molecular targets through which contraction can alter circadian clock phase. Overall, the findings in the current investigation demonstrate that a single bout of exercise or contractions is sufficient to alter the phase of the muscle clock in vivo and in vitro. In addition, the impact of exercise on the muscle clock is dependent on the time of exercise confirming that contractions are a bona fide muscle clock time cue.

The circadian clock has emerged as an important contributor to health and disease (Roenneberg and Merrow, 2016) and the interaction of exercise and the circadian clock has drawn attention (Gabriel and Zierath, 2019; Wolff and Esser, 2012). Studies from as early as the eighties have shown that exercise is a time cue for circadian rhythms as interventions performed in rodents and humans shift the phase of circadian behaviour (locomotion, sleep/wake) depending on time of exercise (Eastman et al., 1995b; Marchant and Mistlberger, 1996; Mistlberger and Skene, 2005; Miyazaki et al., 2001; Mrosovsky, 1996; Reebs and Mrosovsky, 1989; Youngstedt et al., 2019). However, it is important to note that these early studies utilized animal behaviour as a readout for the effect of exercise on phase shifting, whereas we have directly assessed the effects of exercise on the skeletal muscle circadian clock mechanism.

Previous work from our laboratory demonstrated that repeated exercise bouts (i.e. run training) phase-shifted the skeletal muscle clock, suggesting that exercise training could be a zeitgeber for the circadian clock in skeletal muscle. Specifically, we reported that four weeks of exercise training (∼28 exercise bouts) induced a phase advance of the muscle circadian clock (Wolff and Esser, 2012). In contrast, our current investigation is the first to report that a single one hour bout of treadmill exercise induced a significant phase shift of the skeletal muscle circadian clock. Moreover, we performed the acute bout of exercise at three different time points over the day; middle of the rest phase, end of rest phase and middle active phase. We found that there was a time of day specific effect on the muscle clock with exercise induced phase advance, phase delay and exercise in the middle of the active phase not shifting the muscle clock. These findings were exciting as they suggest that exercise is functioning as a true time cue for the circadian clock in skeletal muscle. In fact, since the mice were maintained in normal light:dark conditions these findings indicate that time of day exercise is sufficient to influence the phase of the muscle clock in a mouse with intact central clock and normal light cues.

Interpretation of the effects of exercise on the *in vivo* muscle circadian clock are complicated by the potential confounding influence of the central clock as well as circulating neurohumoral factors (Schibler et al., 2015). Thus, we sought to determine if muscle contractions, as a component of exercise, shift the phase of the muscle circadian clock under constant conditions. The capacity to test a putative time cue under constant conditions strengthens our ability to assess contractions act as a bona fide time cue. We found that like treadmill running, a single one hour bout of electrical stimulation of C2C12 myotubes resulted in either a phase advance or phase delay. Additionally, the phase changes were not simply an acute change to the contractions but rather the phase changes were maintained over days. These maintained changes in phase are in contrast to a local reset, where a stimulus only alters phase for a short duration. Additionally, plotting the effects of contractions on muscle circadian phase (i.e. using the peak of Bmal1 mRNA) reveals a pattern resembling previously reported type 1 phase response curve, with gradual changes between advances and delays (Glass and Winfree, 1984; Winfree, 1977). While these observations indicate that contractions are likely a time cue for the skeletal muscle circadian clock, bona fide time cues must also induce direct changes in the transcriptional machinery of the circadian clock.

The effects of light on the transcriptional response of the core molecular clock in the SCN consistently upregulate *Per1/2* expression, suggesting the negative arm of the molecular clock may be targeted to alter phase (Albrecht et al., 1997; Shigeyoshi et al., 1997; Miyake et al., 2000). In contrast to the effect of light on the SCN, we found that contractions at 22h p.s. and 28h p.s. decreased *Per2* and *Per1/2* mRNA levels, respectively. Our findings are supported by recent data demonstrating that both acute cycling and resistance exercise downregulate *Per1* at four hours post-exercise in humans (Dickinson et al., 2018). However another investigation found that *Per2* was upregulated six hours following an acute bout of resistance exercise (Zambon et al., 2003). This discrepancy may be influenced by the exercise timing, as the participants with a reduction in *Per1* exercised in the morning (Dickinson et al., 2018) and those with an increase in *Per2* exercised in the early afternoon (Zambon et al., 2003). Although these data clearly indicate that the negative arm of the molecular clock are targets of exercise in skeletal muscle, our findings are novel in that we assessed phase changes as well as transcriptional outcomes. Specifically, contractions at 22h p.s. induced a phase advance accompanied by decreased expression of *Per2* mRNA, while contractions at 28h p.s. also lowered the expression of *Per2* mRNA, yet induced a phase delay. We also identified a significant decrease in *Bmal1* expression accompanied by a phase delay at 28h p.s., which is unusual and this may reflect an additional, novel mechanism through which contractions alter skeletal muscle circadian phase.

Finally, while phase changes after contractions at 22h and 28h p.s. were linked to changes in mRNA expression of *Per1/2* and *Bmal1* this was not the case in 40h p.s. cells. Following contractions at 40h p.s. no core clock genes were differentially expressed compared to control despite a phase delay comparable to that at 28h p.s.. Although it is unclear precisely what mechanism was responsible for the phase shift at 40h p.s., previous data have suggested post translational modifications also influence the molecular clock (Blau, 2008; Eng et al., 2017; Noguchi et al., 2018; Sun et al., 2019). Future investigations using the present model system will allow for experiments that can address those questions in depth. Overall, our findings that contractions 1) induce a shift in the phase of the muscle circadian clock in a manner that is similar to a phase 1 Phase Response Curve (PRC), 2) this phase shift is maintained over time and 3) alterations in core clock mRNA expression levels confirms that contractions are a bona fide time cue for the clock in skeletal muscle. In addition, the fact that contractions induce a differential outcome in terms of shifting phase and modification of molecular clock gene expression shows that there is a clear interaction between contractions and the circadian clock in skeletal muscle.

The integration of exercise physiology with circadian biology has revealed the importance of exercise timing on muscle function, metabolism and exercise performance (Gabriel and Zierath, 2019; Tahara et al., 2015; Wolff and Esser, 2019, 2012). In the present study we present evidence defining that muscle contractions are a part of exercise that functions a bona fide time cue for the skeletal muscle circadian clock. This finding could be of interest as exercise has the potential to work as therapeutic to battle negative health outcomes linked to a lifestyle promoting circadian disruption as seen in e.g. shift workers (Jørgensen et al., 2017; Koshy et al., 2019; Morris et al., 2016; Zimmet et al., 2019). Our finding that exercise is a time cue for the skeletal muscle circadian clock in nocturnal rodents as well as in isolated muscle cells that are neither nocturnal or diurnal, indicates that exercise could have similar effects in humans,. Further our newly developed in-vitro system will allow for further studies aiming to elucidate the molecular mechanism that links muscle contractions to the muscle clock by providing a well-defined and isolated system. In the future this could lead to a better understanding of how different time cues work isolated from each other but also in combination which could lead to further improvement of therapeutic strategies combining more than one time to maximize outcome.

## Acknowledgements

The authors are grateful for the technical assistance provided by Collin M. Douglas and constructive discussions with Miguel A. Gutierrez-Monreal. This work was supported by the National Institutes of Health awards R01AR066082 and U01AG055137 to KAE.

## Authors contribution

D.K., C.W and K.E designed the experiments. D.K., C.W. performed the experiments. D.K. C.W analysed the data. D.K, C.W. and K.E wrote the manuscript. All authors reviewed the manuscript and approved the final version. All authors agree be accountable for all aspects of the work in ensuring that questions related to the accuracy or integrity of any part of the work are appropriately investigated and resolved. All persons designated as authors qualify for authorship, and all those who qualify for authorship are listed.

